# Epigenomics of embryogenesis in turbot (*Scophthalmus maximus*)

**DOI:** 10.1101/2024.12.12.628162

**Authors:** Oscar Aramburu, Belén G. Pardo, Ada Jimenez-Gonzalez, Andrés Blanco-Hortas, Daniel J. Macqueen, Carmen Bouza, Paulino Martínez

## Abstract

Embryogenesis is the crucial first step of ontogeny, where an organism with a complex body plan arises from a single undifferentiated totipotent cell. This process is orchestrated by dynamic changes in transcriptional regulation, influenced by chromatin accessibility and nucleotide and histone modifications constituting epigenetic signals enabling access to transcription factors. The epigenomic regulation of embryogenesis has been studied in model fishes, but little attention has been paid to farmed fish - where traits of importance to aquaculture rely on early developmental processes. This study, framed within the AQUA-FAANG consortium, reports a comprehensive regulatory atlas of embryogenesis for turbot (*Scophthalmus maximus*), a farmed flatfish representing order Pleuronectiformes. 14,560 genes were expressed in the embryonic transcriptome with > 90% showing differential expression across consecutive stages. By integrating multi-histone ChIP-Seq marks with ATAC-Seq, we built a genome-wide chromatin state model, defining promoter and enhancer activity across stages. Transcription factor binding motif (TFBM) analysis of differentially active promoters and enhancers revealed dynamism in regulated gene functions, with more than half the TFBM enriched in a single developmental transition. Significant shifts in chromatin accessibility occurred across stages, most notably during the transition from shield to early segmentation, suggesting a profound chromatin reorganization underpins somitogenesis and early organ development. Most changes in chromatin accessibility across stages did not involve promoter regions of differentially expressed genes, suggesting a trend of promoter accessibility preceding gene transcriptional activity. Comparative analyses with zebrafish revealed a global transcriptomic correlation of single copy orthologs at matched stages of embryogenesis across species. While conserved expression dynamics were revealed for many orthologous Hox genes, notable cross-species differences were identified from before zygotic genome activation leading up to hatching. This multi-omics investigation provides a novel atlas of non-coding regulatory elements controlling turbot development, with key applications for flatfish biology and enhancing sustainable aquaculture.

## INTRODUCTION

Embryogenesis is a fundamental process in the life cycle of multicellular organisms. Insights into embryonic development can shed light on biological questions such as its genetic control, inter-species variation and evolution. Research on embryogenesis is not only limited to basic biological knowledge, but also crucial for understanding genetic and epigenetic mechanisms behind the establishment of cell lineages and tissue formation (Petratou et al., 2024). Such insights can help optimize breeding strategies and improve the management of commercially valuable species farmed in controlled environments.

Embryogenesis involves a dynamic succession of key processes such as cell differentiation, migration, patterning, organogenesis and apoptosis (Zernicka-Goetz, 2002; Rasmussen, 2003; Kurosaka and Kashina, 2009; Voss and Strasser, 2020). These processes are particularly sensitive to environmental stressors such as pH, salinity, pathogens and temperature (dos Santos et al., 2020; Feugere et al., 2021; Miranda et al., 2023), currently one of the most acute issues in aquaculture due to global warming (Skjærven et al., 2024). By understanding gene regulatory interactions underlying these processes, genetic variants linked to favourable traits can be identified and targeted by breeders.

Annotation of the genome of aquaculture species represents an essential resource for understanding the genetic and functional genomic mechanisms underlying complex traits such as growth, feed efficiency, muscle quality, reproduction, and disease resistance (Clark et al., 2020; Houston et al., 2020; Johnston et al., 2024). Efforts in functional annotation of finfish genomes have to date concentrated on coding and non-coding genes, with transcriptomic studies establishing detailed annotations in zebrafish (Presslauer et al., 2017; White et al., 2017; Lawson et al., 2020) and aquaculture species including Atlantic salmon (Bizuayehu et al., 2019; Gervais et al., 2022; Ignatz et al., 2022; Shwe et al., 2022; Abdellaoui and Kim, 2024), rainbow trout (Ma et al., 2019; Cleveland et al., 2020; Yang et al., 2023), Nile tilapia (Powell et al., 2021; Kumar Behera et al., 2024) and European sea bass (Lama et al., 2020; Faggion et al., 2021; Papadaki et al., 2022), among others (Li et al., 2021; Blasweiler et al., 2023; Wang et al., 2023; Bouza et al., 2024; Duarte-Ribeiro et al., 2024; Villareal et al., 2024).

In contrast, *cis*-regulatory elements (CRE), regulatory sequences including promoters, enhancers, silencers, and insulators, remain largely unexplored in aquaculture species (Song et al., 2022; Harvey et al., 2024), despite recent advances made in model species such as zebrafish (Yang et al., 2020; Baranasic et al., 2022; Jimenez-Gonzalez et al., 2023) and terrestrial livestock (Summers et al., 2020; Pan et al., 2021; 2023; Xiang et al., 2021). These elements are crucial for regulating gene expression, and their activity can vary depending on ontogeny, tissue, cell, sex, age, and health status of the organism (Kellis et al., 2014; van Mierlo et al., 2023). Therefore, annotation of these elements will help address fundamental biological questions on development and tissue differentiation (White et al., 2017; Halstead et al., 2020; Pan et al., 2023), but also understanding phenotypic plasticity in response to environmental changes (Hu and Barrett, 2017; Liu et al., 2023; Bouza et al., 2024; Hu et al., 2024).

Turbot (*Scophthalmus maximus*) is a valued fish species farmed primarily in Asia and Europe, with an annual production exceeding 100,000 tons largely concentrated in China (Gao et al., 2023), followed by Spain (APROMAR 2023). European turbot breeding is in its eight generation of selection (Riaza, Stolt Sea Farm SA, pers. comm.). Functional annotation of the turbot transcriptome has been conducted using high-quality genome assemblies (Figueras et al., 2016; Maroso et al., 2018; Xu et al., 2020; 2022; Martínez et al., 2021), but non-coding elements have received limited attention, aside from initial studies exploring non-coding RNAs (Robledo et al., 2017; Xue et al., 2021; Cai et al., 2022; 2023). The morphological processes underlying the transformation of a fertilised egg into a free-living flatfish are well-established in turbot (Jones, 1972; Devauchelle et al., 1988; Pepe-Victoriano et al., 2012; Tong et al., 2012; Wang et al., 2017). However, few studies have focused on underpinning developmental changes in gene expression (Wu et al., 2024), with only one investigating regulation of chromatin accessibility during flatfish metamorphosis, using turbot as a study system (Guerrero-Peña et al., 2023).

In this study, using a highly contiguous chromosome-level genome assembly (Martínez et al., 2021), we generated the first comprehensive functional annotation of turbot embryogenesis. RNA-Seq, ATAC-Seq and ChIP-Seq was performed on replicated embryo samples from different developmental stages to assess integrated changes in the transcriptome, chromatin accessibility, and epigenetic states determined using functionally distinct (H3K4me3, H3K27ac and H3K27me3) histone marks. Comparative transcriptomics of turbot against zebrafish embryogenesis data (White et al., 2017) was performed with a focus on the Hox gene family, which includes key regulators of anterior-posterior body plan during embryogenesis., Although highly conserved among vertebrates, divergence in Hox family has been linked to evolutionary innovations contributing to body structure differences across species (Cumplido et al., 2024). Our findings comprehensively advance understanding of the epigenomic mechanisms underlying turbot embryogenesis in comparison to a model fish species, providing a new biological resource for prioritizing genetic variants linked to regulatory elements controlling traits shaped during early development relevant for breeding programs.

## RESULTS

114 multiomic datasets were produced in this study, including RNA-Seq (36), ATAC-Seq (18) and ChIP-Seq (49; 17 for H3K4me3, 17 for H3K27ac and 15 for H3K27me3, plus 6 ChIP-Seq input controls). Full information on samples and metadata is shown in **Supplementary tables 1 and 2**.

### RNA-Seq

On average, 64,550,207 raw reads per library were produced across the 36 RNA-Seq libraries, with 97.59% mapping rate to the turbot genome (range: 97.21-98.20; **Supplementary table 3A**). Principal component analysis (PCA) separated all 12 developmental stages following a smooth ellipsis from the cleavage stage (64 cells) up to the end of embryogenesis (prehatch). For all stages, the three replicates clustered closely in the PCA and the heatmap visualising pairwise sample expression similarity (**Figure 1A and B**). Moreover, the hierarchical clustering heatmap grouped developmental stages in three major clusters, separated by the zygotic genome activation (ZGA): pre-ZGA (64-cells, morula and early blastula); ZGA (late blastula and embryonic shield stages); and post-ZGA (neural plate, blastopore closure, early, mid and late segmentations, early pharyngula and prehatch). Further subdivision was detected, particularly for the two stages at ZGA (late blastula and shield), and post-ZGA (before and after mid segmentation; **Figure 1B**).

**Figure 1.**
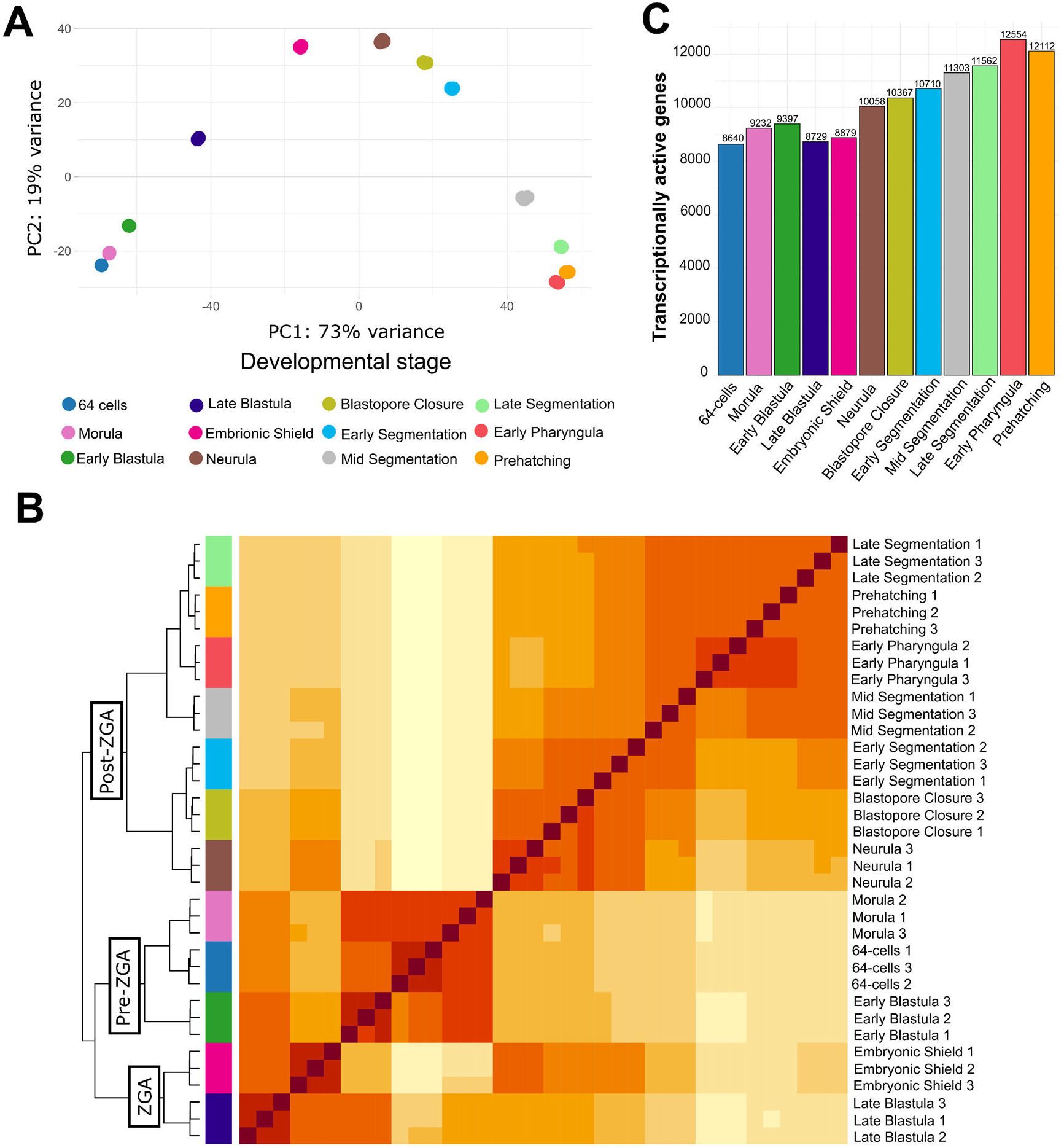
**A)** PCA and **B)** heatmap (Spearman correlation) of RNA-Seq data on the 12 developmental stages and replicates studied; **C)** number of active genes (TPM > 5) for each developmental stage.

14,560 total expressed genes (TPM > 5 cut-off; representing 42.51 % of 34,252 annotated genes) were identified, constituting the developmental transcriptome of turbot (**Supplementary table 4**). A trend of increasing gene expression was observed from early (e.g. 8,640 genes at 64-cell stage) to late (e.g. 12,112 genes during prehatch stage) stages (**Figure 1C**). Correlation networks using BioLayout (**Figure 2A**; Theocharidis et al., 2009) were applied to visualise the transition of expression across stages, ordered temporarily by stage of maximum expression in 12 clusters (**Figure 2B**). The correlation network shows a “pulley-arch” structure, where the clusters on the left- and right-hand side of the graph represent genes expressed at early or late stages, respectively. Both groups are not completely isolated but joined by genes that reached peaks of expression during gastrulation and segmentation stages, as well as by a small set of genes with high expression at early and late stages along the “string” of the bow-shaped graph (**Figure 2A**). Detailed profiles for a selection of developmental genes associated with maternal expression, ZGA, writing and reading of epigenetic marks, cellular differentiation and organogenesis can be found in **Supplementary figure 1**, showing the expected expression patterns based on the literature (see discussion section).

**Figure 2.**
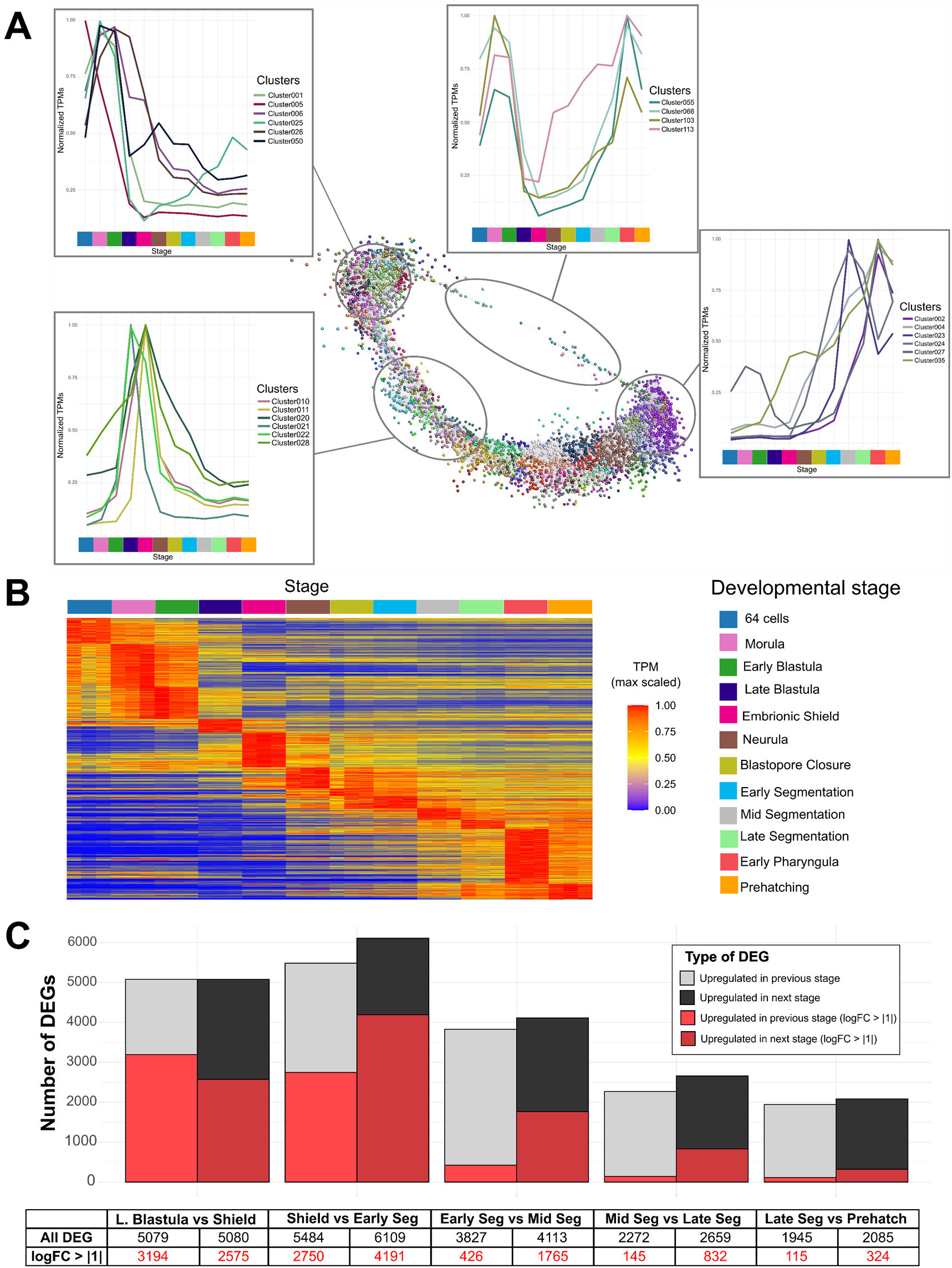
**A)** BioLayout correlation networks showing the transition of expression patterns across developmental stages; **B)** Heatmap of the expression profiles across developmental stages. Expression values are scaled to the maximum expression for each gene, across development, and genes are organised by the stage where the maximum expression occurs, then clustered within each stage. **C)** Number of differentially expressed genes (DEGs) for each comparison between consecutive developmental stages. In red, number of DEGs with fold change (FC) > 1.

Differential expression analysis was performed by comparing the six main developmental stages sequentially, resulting in five distinct comparisons. In total, 14,060 differentially expressed genes (DEG; p.adj < 0.05) were identified across between-stage comparisons, representing 96.6% of the total active transcriptome (**Figure 2C**; **Supplementary table 5**). Overall, more DEGs were identified at early and mid-development (late blastula to shield transition; early to mid-segmentation transition) compared to later developmental stages (**Figure 2C**; **Supplementary table 5; Supplementary figure 2**). For each comparison, similar proportions of DEGs were found in the two compared stages with more extreme DEGs (logFC > 1.0) usually more frequently upregulated at the “next” stage (**Figure 2C**).

### High-resolution functional enrichment profiles of turbot embryogenesis

Gene Ontology (GO) analysis was used to identify enriched terms associated with biological processes (BP) for the following lists of genes (**Supplementary table 6**): i) the total active transcriptome (**Figure 3A**), ii) total inactive transcriptome (**Supplementary figure 3**), iii) stage-specific active transcriptomes (12 stages; **Supplementary figure 3**), and iv) the DEGs between the five sequential-stage comparisons (**Figure 3B; Supplementary figure 3**).

**Figure 3.**
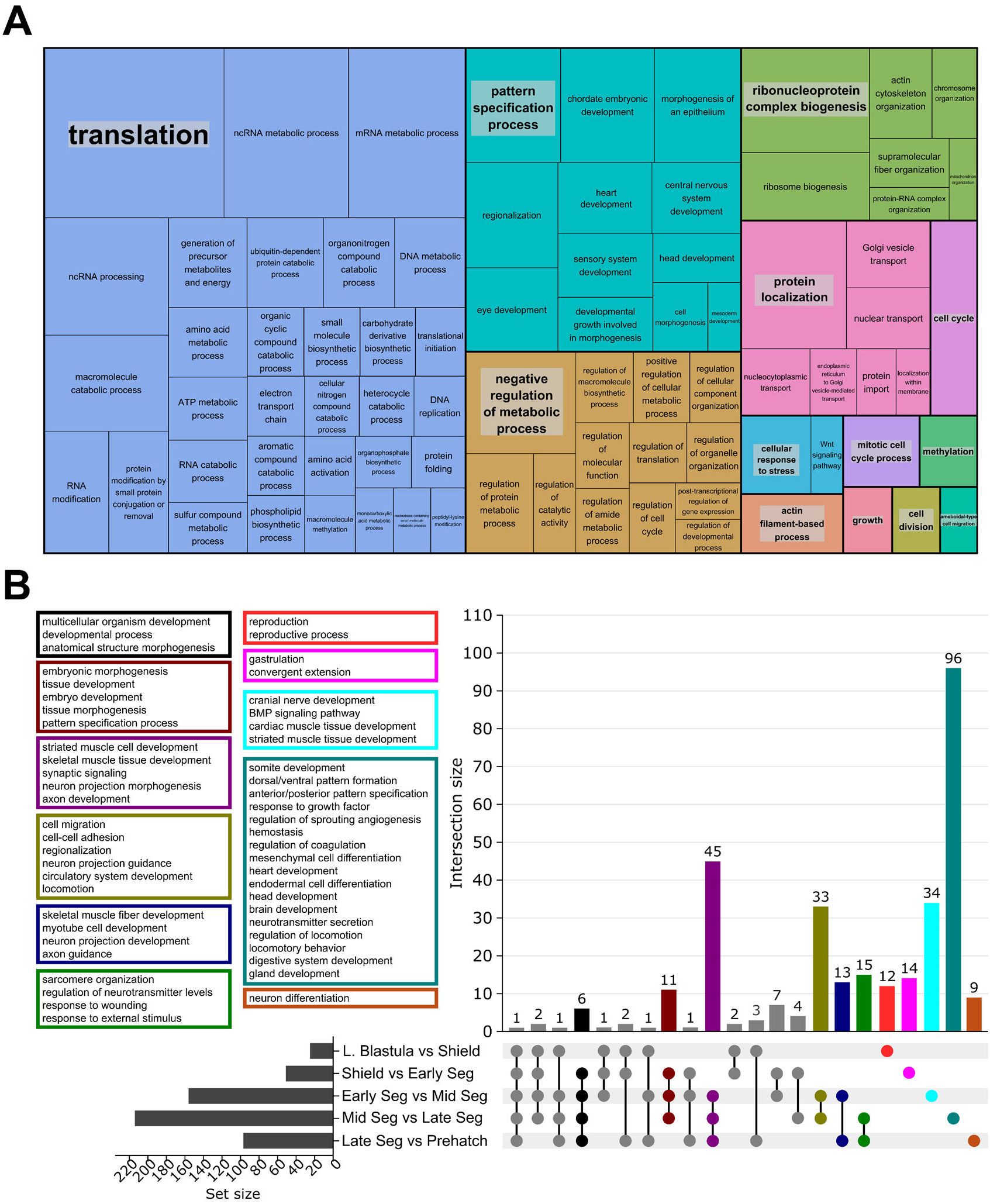
**A)** Treemap chart of enriched GO terms (Biological Process, BP) detected in the total active transcriptome of turbot during embryogenesis; terms related to similar functions are grouped by the same colour and the area of each term is inversely proportional to its FDR p-value. **B)** UpSet plot showing the overlapping enriched GO terms between consecutive developmental stages; each coloured box corresponds to the same coloured barplot, summarizing BP terms.

The most prominent groups of enriched GO terms in the total active transcriptome were associated with metabolism, replication, transcription and translation processes, followed by terms associated with pattern specification processes and organogenesis (**Figure 3A; Supplementary table 6**). A similar pattern was observed through all the 12 developmental stages (**Supplementary figure 2**), where terms related to transcription, translation, ribosome biogenesis and protein localization were shared among all stages. However, some terms were specific to certain stages, including cell cycle regulation, detected across early developmental stages (especially from late blastula to early segmentation); reproductive process at the cleavage / early blastulation stages (64 cell, morula and early blastula); methylation-related terms (RNA, protein and macromolecule methylation) at all developmental stages except from late segmentation onwards; endoderm formation in late blastula and embryonic shield stages; and morphogenesis-related terms in early segmentation (mesenchymal tissues, epithelium and glial cells), late segmentation (actin organization), and early pharyngula (neuron differentiation) (**Figure 3A**, **Supplementary figure 3, Supplementary table 6**).

For the total inactive transcriptome (19,893 annotated genes), many enriched GO terms were associated with immune processes, response to stimulus, presynapse organization, regulated exocytosis, feeding behaviour, and cell adhesion / communication (**Supplementary Figure 3; Supplementary table 6**).

RNA processing and transcription associated terms were enriched across all the five pairwise stage comparisons, suggesting a fine adjustment of these processes all along development (**Figure 3B; Supplementary table 6**). More stage-specific processes, including reproductive processes, were found only in the late blastula to shield transition. Inspection of these genes revealed a variety of reproductive and germline markers of oocyte (*zp3b*, *bmp15*, *figla*) and sperm development (*spata6*, *spaca4l*, *pgrmc1/2*), upregulated in late blastula, and different stem cell and pluripotency factors (*nanog*, *kdm6a*, *pum1*, *pum3*) upregulated towards shield stage. Moreover, gastrulation, convergent extension, pattern specification and tissue (epithelium) morphogenesis in the transition from shield to early segmentation; morphogenesis-related GO terms (particularly muscle development and nerve development/synapsis) in all the transitions across segmentation, while head, heart, blood, digestive system and gland development-related terms were very prominent in mid to late segmentation; and terms related with locomotion and behaviour were enriched in the transitions from both mid to late segmentation and late segmentation to prehatch, the latter also showing terms associated with response to stimulus (**Figure 3B; Supplementary Figure 3; Supplementary table 6**).

### ATAC-Seq and ChIP-Seq

After sequencing, 75,600,169 (ATAC-Seq), 113,393,477 (H3K27ac), 98,992,315 (H3K27me3) and 121,913,111 (H3K4me3) raw reads were produced on average, with a mean mapping rate of 93.54% to the turbot genome (**Supplementary table 3B**). On average, 50,004 peaks were identified for ATAC-Seq libraries across all stages, while 18,466 (H3K4me3), 12,127 (H3K27ac) and 16,651 (H3K27me3) peaks were called for the ChIP-Seq data (**Figure 4A**). Hierarchical clustering (Spearman correlation) clustered each library type, with some overlap between H3K4me3 and H3K27ac (**Figure 4B**). ATAC-Seq and ChIP-Seq libraries were mostly segregated across PC1 (35%), while PC2 (19%) explained most of the variance between histone marks, particularly differentiating the repressor mark H3K27me3 from the nearby activation marks H3K4me3 and H3K27ac (**Figures 4B and C**). All four techniques differentiated clearly the ZGA stages (late blastula and shield) from post-ZGA stages, and ATAC-Seq showed the greatest resolution to differentiate each developmental stage (**Figure 4C**).

**Figure 4.**
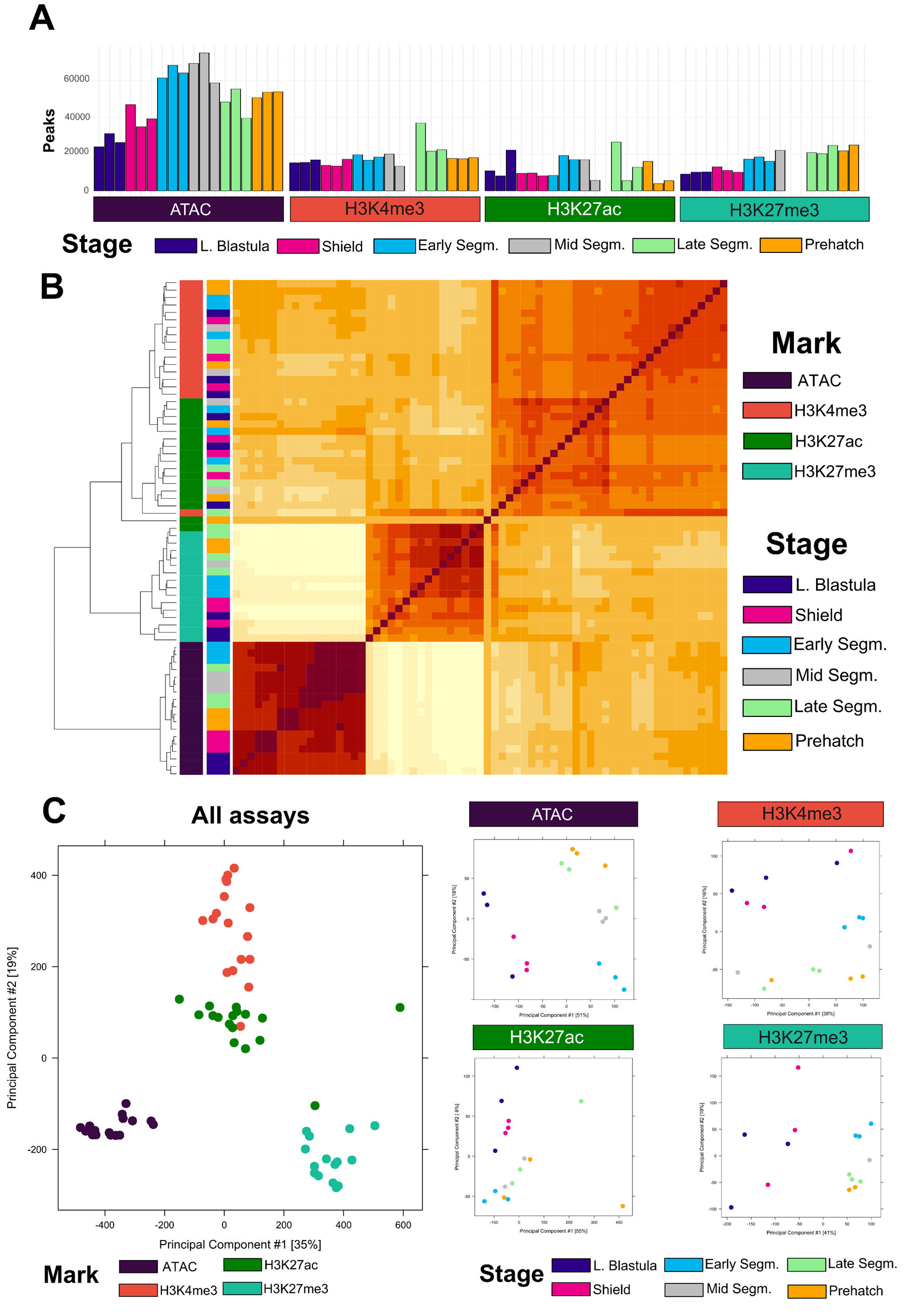
**A)** Number of peaks called for each chromatin assay, developmental stage, and replicate; **B)** Heatmap (Spearman correlation); and **C)** PCA of ATAC-Seq and ChIP-Seq data on the 6 main developmental stages studied.

### Chromatin state annotation

Genome-wide chromatin state predictions were obtained by integrating the ChIP-Seq and ATAC-Seq datasets across stages using ChromHMM (**Figure 5A**, **Supplementary table 7**). After iterations at different state counts, a 10-state model was chosen as it maximized the number of non-redundant biologically relevant states. This model included four active promoter states (States 1 to 4), one active enhancer state (7), one weak active regulatory element without ATAC (5), one bivalent regulatory element state (6), one ATAC island state (8), one Polycomb state (9) and one low signal state (10). States 1 to 5 (active promoter states and weak active regulatory element without ATAC) exhibited the highest signal around the TSS (±2 kb), corresponding to promoter regions and/or transcriptionally active regions.

**Figure 5.**
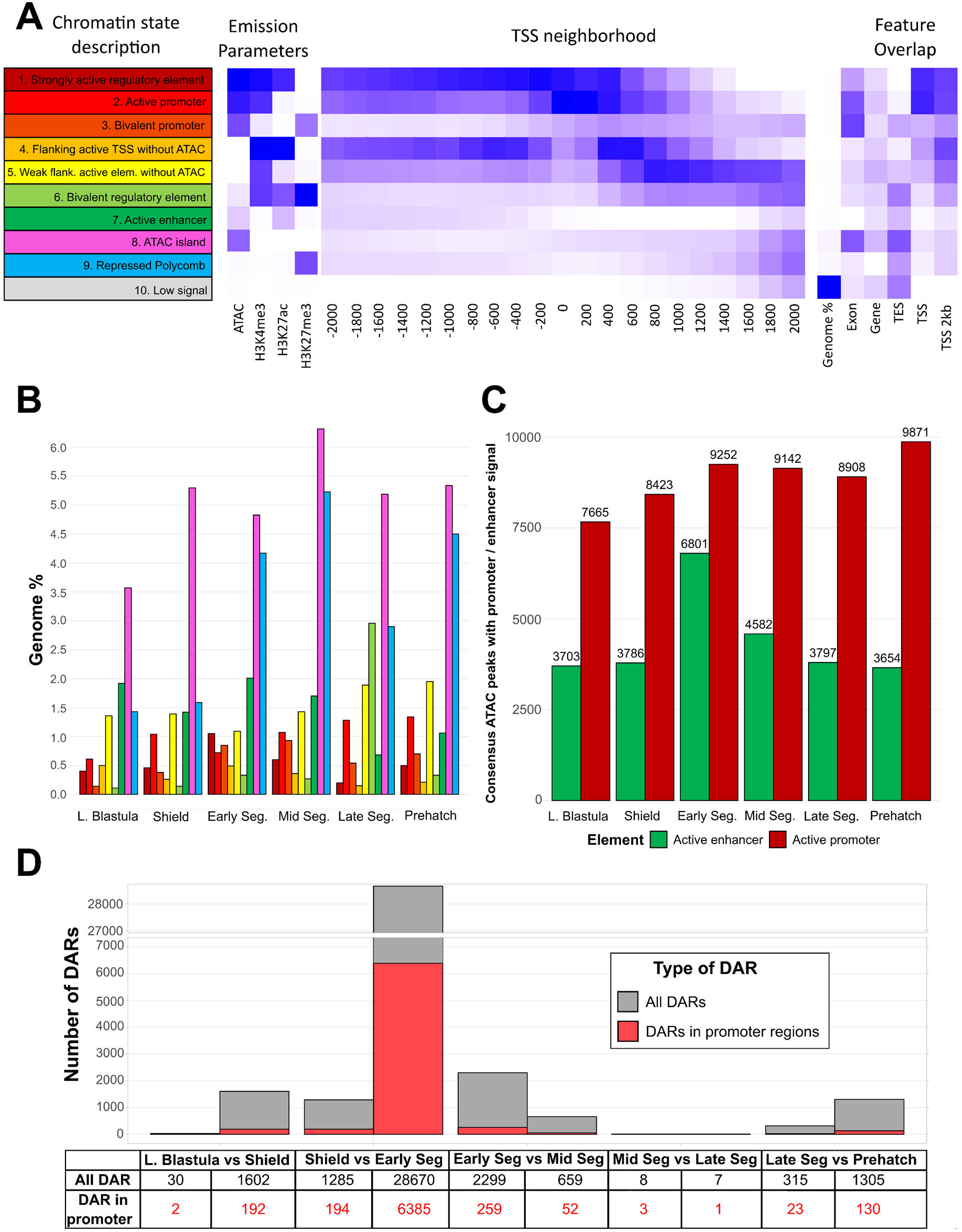
**A)** 10-state chromatin model of the turbot genome during embryogenesis: heatmaps of the emission parameters for each chromatin state are shown for ChIP-Seq data with three histone marks (H3K4me3, H3K27ac and H3K27me3) and ATAC-Seq (left), the emission parameters of each chromatin state considered in the neighbourhood around the TSS (middle) and alongside the features in the turbot genome (right). TSS: Transcription start site; TES: Transcription end site. **B)** Average genomic coverage (%) per developmental stage for each chromatin state (Low signal state excluded). **C)** Consensus ATAC-Seq peaks overlapping active promoter / enhancer chromatin states (states 1 to 7). **D)** Number of DAR for each comparison between consecutive developmental stages. In red, number of DARs located within promoters.

The low signal state (state 10) covered most of the turbot genome (85.47% on average). The ATAC island (state 8) was the most prominent open chromatin state (5.09%), while active promoter (1 to 4) and enhancer (7) states represented 2.46% and 1.46% of the genome sequence, respectively. Unspecific weak active and bivalent states (5 and 6, respectively) represented 2.21% of the genome, bringing the total coverage of accessible, activation-marked regions to 6.13% (**Figure 5B)**. Finally, the repressed Polycomb state (state 9) marked by H3K27me3 averaged 3.30%.

Consensus ATAC-Seq peaks overlapping promoter regions (-2kb, +600bp) and intergenic / intron regions (putative enhancers) were intersected with the annotated chromatin states to determine active CREs in each condition (**Figure 5C; Supplementary table 8**). On average, 14,934 open chromatin regions with active regulatory signals were found across the six stages, ranging from 13,217 in late blastula to 18,136 in early segmentation. Promoter activity showed a steady increase from late blastula (7,746) to prehatch (9,988), while enhancer activity showed a rapid increase from shield (5,237) to early segmentation (8,832), decaying along mid segmentation (6,231) to reach similar levels to early development (average 5,205; **Figure 5C**). Detailed chromatin profiles for relevant genes associated with maternal expression, ZGA, writing and reading of histone marks, cellular differentiation and organogenesis are presented in **Supplementary figure 4**, showing a generally conserved accessibility of the promoter through the analysed developmental stages.

### Differentially accessible regions and intersection with DEGs

Significantly differentially accessible regions (DARs, FDR-adjusted *p* < 0.05) between consecutive developmental stages were identified (**Figure 5D**, **Supplementary figure 5**; **Supplementary table 9**). By far, the highest number of DARs were detected in the transition from embryonic shield to early segmentation (28,686 DARs), while only 14 DARs were found in the transition from mid to late segmentation. On average 12.64% DARs were found within promoter regions and 20.86% in intergenic and intronic regions (**Figure 5D**; **Supplementary table 9**).

The intersection between DARs and DEGs for each comparison was tested through hypergeometric distribution tests (p.adj < 0.05, Bonferroni correction, **Supplementary table 10**). Little correspondence was found between DARs in promoters and DEGs, with the exception of the transition from shield to early segmentation, with 2,034 DEGs with DAR promoters upregulated in early segmentation (p-value 6.48e-08; 33.29 % of the DEGs and 35.89 % of the promoter-DARs in that transition) and the transition from late segmentation to prehatch, with 42 DEGs with DAR promoters upregulated in prehatch (p-value 1.86e-10, 2.01 % of the DEGs and 36.52 % of the promoter-DARs in that transition), suggesting that generally, promoters of DEGs are already accessible before the onset of transcription.

### Transcription factor binding motif analysis

To identify the main TFs associated with chromatin accessibility dynamics in candidate CREs regulated during embryogenesis, TFBM enrichment was conducted for all DARs associated with promoters and enhancers. Significant TFBM enrichments were observed for all transitions, except from mid to late segmentation (**Figure 6A; Supplementary table 11**). The highest number of significantly enriched TFBM was detected in the shield to early segmentation transition (74 motifs) followed by early to mid-segmentation (37), late blastula to shield (19) and late segmentation to prehatch (5).

**Figure 6.**
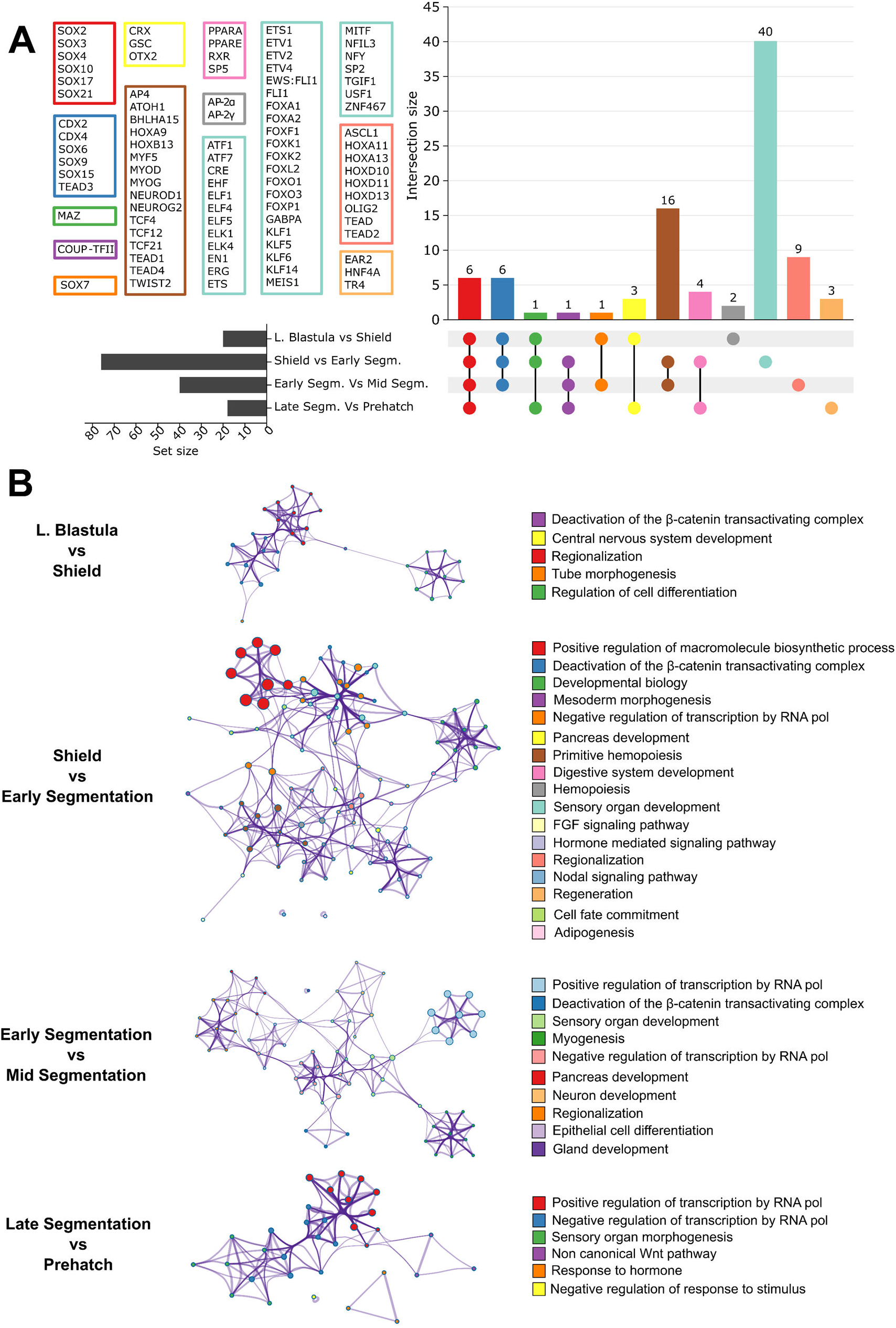
Enrichment of transcription factor (TF) binding motifs (TFBM) within promoter and enhancer elements of the turbot genome showing differentially chromatin accessibility between consecutive developmental stages. **A)** UpSet plot of enriched TFBM shared between comparisons; each coloured box corresponds to the same-coloured histogram bar. **B**) Ontology enrichment clusters of TFs explaining each enriched TFBM; each cluster is coloured with the most statistically significant term among terms clustered together; the size of each term is given by -log_10_p and the stronger the similarity among nearby terms, the thicker the edges between them.

Out of the 92 enriched motifs identified among DARs, 12 were enriched in all transitions spanning the late blastula to mid segmentation stages; 8 were identified as binding motifs to different SOX family TFs (**Figure 6A; Supplementary table 11**). 16 TFBM were enriched in transitions from shield to early segmentation and early to mid-segmentation. These represented TFs involved in muscle (MYOD) and neural (NEUROD1, NEUROG2) differentiation, tissue growth and cell proliferation (TEAD1, TEAD4) and two Hox proteins (HOXA9 and HOXB13), although most of the HOX binding motifs detected (five) were exclusive to the early to mid-segmentation transition (along with TEAD motifs). Finally, 40 binding motifs were exclusively enriched in the shield to early segmentation transition. This group of TFBM mainly comprised members of the ETS and FOX TF families, two of the largest families of transcriptional regulators in metazoans, involved in cell proliferation, differentiation and development, as well as immune response (**Figure 6A; Supplementary table 11**).

To further investigate the functions of TFs involved in turbot embryogenesis, the molecular Complex Detection algorithm (MCODE) was applied to identify clustered functions associated with TFs explaining all enriched TFBM (**Figure 6B**). The only enriched function detected through all transitions was the deactivation of the β-catenin transactivating component (involved in Wnt signaling pathway). Clustering for the late blastula to shield transitions unveiled functions related to regionalization, cell differentiation and development of the central nervous system, endocrine system and inner ear. During the shield to early segmentation transition, more organ / tissue development functions arise (primitive hemopoiesis, epithelial and muscle differentiation, heart development, mesoderm development, adipogenesis) that continue towards the early to mid-segmentation transition with the addition of other organ development functions (sensory organ, pancreas and digestive system development). No clustering was detected for the late segmentation to prehatch transition.

### Comparative developmental transcriptomics

The developmental transcriptome of turbot and zebrafish was compared considering the six stages profiled for turbot (late blastula, embryonic shield, early segmentation, mid segmentation, late segmentation and prehatch) versus matched stages from zebrafish defined by White et al. (2017): dome, gastrula shield, 1-4 somites, 14-19 somites, 20-24 somites and long pec, respectively.

14,221 orthogroups were established between turbot and zebrafish; 8,466 genes constituted single copy orthogroups, with 6,967 shared genes showing expression (cut-off TPM > 5) in any of the six stages (**Supplementary figure 6**; **Supplementary table 12**). However, some of these genes showed expression only in one species, 658 for turbot and 817 for zebrafish, with 5,491 expressed in both species (**Supplementary figure 6**; **Supplementary table 12**).

Hierarchical clusteringafter inter-species normalization of the expressed orthologs for each species (Spearman correlation) showed moderate correlations between expression patterns, with some genes showing varying levels of asynchrony between both species (**Supplementary figure 6**). Enriched GO terms were only detected in the exclusive-zebrafish genes, with most functions associated with early development functions, like reproductive processes, gamete generation, cell signalling and adhesion (**Supplementary figure 6**; **Supplementary table 12**). Although no significant GO enrichment was detected in the exclusive-turbot genes, some notable genes were identified such as dystrophin (*dmd*; muscle development), patched 1(*ptch1*; hedgehog signaling pathway), GATA binding protein 4 (*gata4;* heart and gut development), KIT proto-oncogene (*kita;* germ and blood cell development), nuclear factor interleukin 3 regulated member 6 (*nfil3-6;* immune system development), cysteine rich transmembrane BMP regulator 1 (*crim1;* blood and neural tissues) and interleukin 1β (*Il-1b*; bone development).

### Comparative analysis of the Hox gene family

To investigate the evolutionary conservation and divergence of Hox genes, key regulators of anterior-posterior axis patterning and segmental identity, we compared the expression profiles of the 46 and 47 annotated Hox family genes annotated in the turbot and zebrafish genomes, respectively. Each species harbours seven colinearly expressed Hox clusters distributed in chromosomes 1, 11, 14, 15, 18, 19 and 22 in turbot and chromosomes 3, 9, 11, 12, 16, 19 and 23 in zebrafish, for a total of 55 Hox genes (38 orthologs, 8 and 9 exclusives to turbot and zebrafish, respectively). HoxCb and HoxDb clusters were missing in turbot and zebrafish, respectively, while HoxBa, HoxBb and HoxD were partially missing in turbot, and HoxAa and HoxBb in zebrafish (**Figure 7; Supplementary table 13**).

**Figure 7.**
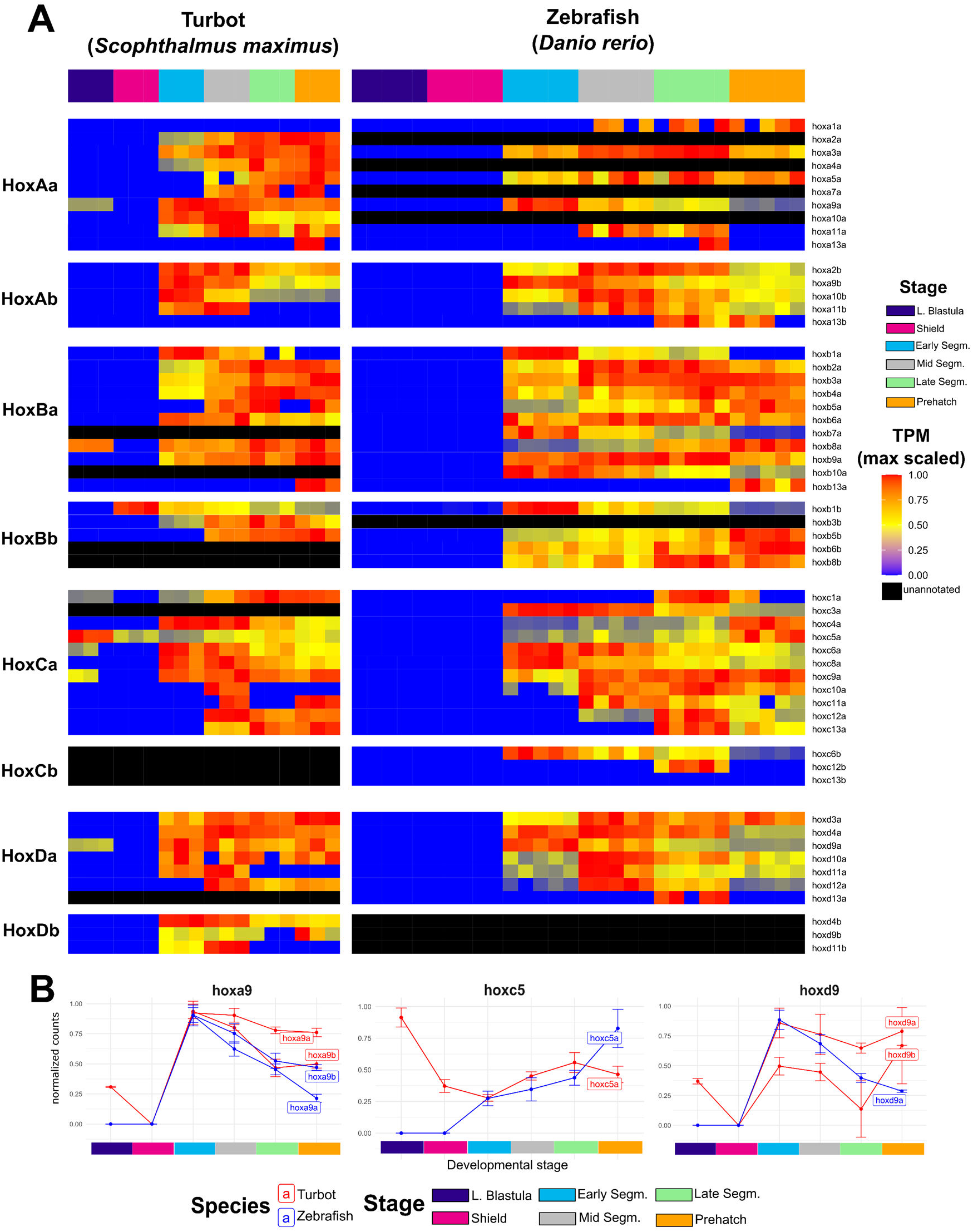
**A**) Heatmap of normalized expression profiles of Hox family genes across 6 matched developmental stages in turbot and zebrafish. **B**) Examples of normalized expression profiles for Hox family genes with i) similar expression between orthologs and paralogs (*hoxa9*), ii) different pattern of orthologs (*hoxc5*), and iii) different pattern between orthologs and paralogs (*hoxd9*).

Comparing the expression patterns between orthologs from zebrafish and turbot, 24 Hox family genes shared very similar expression patterns between both species (**Table 1**; **Figure 7; Supplementary table 13; Supplementary Figure 7**). Eight, however, showed variable degrees of asynchrony, for example, with *hoxa2b* in zebrafish reaching its peak of expression during mid-segmentation instead of early segmentation in turbot. Twelve Hox genes showed strikingly different expression patterns. Two were not expressed in turbot (although the gene is annotated) and the remaining 17 *hox* genes were not present in the genome of either species.

**Table 1.** Categorization of Hox family genes based on the expression pattern similarity between turbot and zebrafish orthologs.

| Category | Number of *hox* genes | *hox* genes |
| --- | --- | --- |
| Similar pattern | 16 | *hoxa3a, hoxa9a, hoxa9b, hoxb2a, hoxb3a, hoxb4a, hoxb6a, hoxb8a, hoxb9a, hoxb13a, hoxc6a, hoxc8a, hoxc9a, hoxd3a, hoxd4a, hoxd12a* |
| Similar but asynchronous pattern | 8 | *hoxa2b, hoxa5a, hoxa10b, hoxb1a, hoxb1b, hoxc12a, hoxc13a, hoxd10a* |
| Different pattern | 12 | *hoxa11a, hoxa11b, hoxa13a, hoxb5a, hoxb5a, hoxc1a, hoxc4a, hoxc5a, hoxc10a, hoxc11a, hoxd9a, hoxd11a* |
| Not expressed in turbot | 2 | *hoxa1a, hoxa13b* |
| Not in turbot genome | 9 | *hoxb6b, hoxb7a, hoxb8b, hoxb10a, hoxc3a, hoxc6b, hoxc12b, hoxc13b, hoxd13a* |
| Not in zebrafish genome | 8 | *hoxa2a, hoxa4a, hoxa7a, hoxa10a, hoxb3b, hoxd4b, hoxd9b, hoxd11b* |

Regarding turbot paralogs (ohnologs), 16 Hox family genes showed similar expression patterns between paralogs. Among them, 6 were asynchronous and 4 showed different expression levels between paralogs (**Table 2**; **Figure 7; Supplementary table 13; Supplementary Figure 7; Supplementary Figure 8**)

**Table 2.**
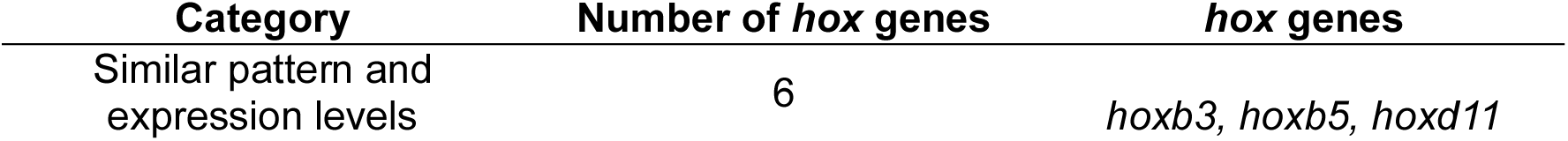
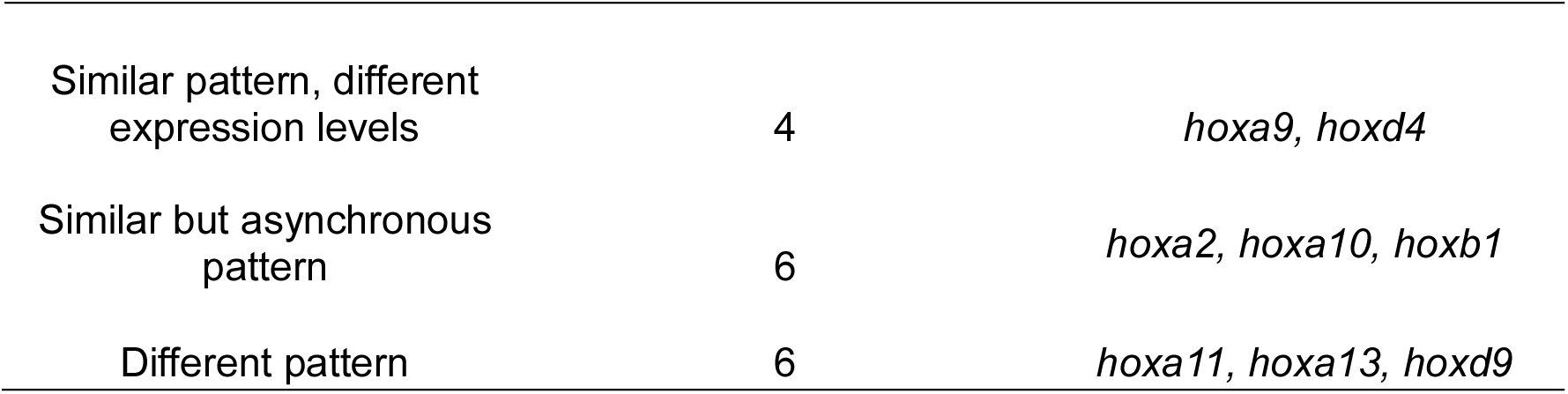
Categorization of Hox family genes based on the expression pattern and expression level similarities between turbot paralogs.

| Category | Number of *hox* genes | *hox* genes |
| --- | --- | --- |
| Similar pattern and expression levels | 6 | *hoxb3, hoxb5, hoxd11* |
| Similar pattern, different expression levels | 4 | *hoxa9, hoxd4* |
| Similar but asynchronous pattern | 6 | *hoxa2, hoxa10, hoxb1* |
| Different pattern | 6 | *hoxa11, hoxa13, hoxd9* |

## DISCUSION

In this study, a multiomic approach capturing epigenomic, transcriptomic and chromatin accessibility dynamics, was used to provide insights into the genetic control of turbot embryo development. Our study also provides a biological resource to understand the genetic basis of traits relevant to sustainable turbot aquaculture.

The transcriptomic analysis showed that most of the active genes and associated GO terms detected across turbot embryogenesis were differentially expressed in the transitions across consecutive developmental, showing the highly dynamic regulation undergoing embryogenesis to specify the fate of the different cell types of the organism (Schwanhäusser et al., 2011; Briggs et al., 2018).

High-resolution chromatin state analyses provided detailed data on the accessibility and epigenetic marks in CREs that regulate developmental processes taking place after ZGA (Fu et al., 2021; Minnoye et al., 2021).

The 10-state chromatin model resulted in the definition in the turbot genome of a variety of chromatin states associated with active promoter and enhancer states, repressed Polycomb regions and ATAC islands, annotated in other species following the same approach (Heintzman et al., 2007; Ernst and Kellis, 2017; Blackledge and Klose, 2021; Barral and Déjardin, 2023), representing the first public annotation of chromatin states during turbot embryogenesis. Interestingly, two of the unveiled chromatin states had a combination of repressor and activation signals, classifying them as bivalent / poised states, which are believed to keep genes ready for expression in response to regulatory cues during embryo development and immune responses (El-Dahr and Saifudeen, 2019; Herrera-Uribe et al., 2020)

### Early development: cleavage, blastulation and the ZGA

At the onset of embryogenesis, during the first synchronised mitotic divisions the embryo is transcriptionally quiescent, and progress is mainly driven by maternal transcripts (Kimmel et al., 1995; Xu et al., 2024). Transcriptomic hierarchical clustering depicted three main developmental stages around ZGA activation, with the cluster of ZGA-stages (late blastula and shield) separating the pre- and post-ZGA stages.

Moreover, all chromatin assays showed similar clustering with ZGA stages well differentiated from post-ZGA stages, particularly noticeable in ATAC-Seq, H3K4me3- and H3K27me3-ChIP-Seq peaks, an expected observation since ZGA triggers the initiation of transcription, where major chromatin remodelling events take place (Schulz and Harrison, 2019). The punctual appearance of GO terms associated with cell cycle regulation, reproductive process and methylation in early developmental stages but not in later ones, is consistent with the presence of parental transcripts, ungapped cell division and highlights the importance of methylation dynamics during early epigenetic reprogramming (Harvey et al., 2013; Messerschmidt et al., 2014; Brantley and di Talia, 2021).

ZGA begins when a specialized set of TFs called pioneer factors (such as those coded by *pou5f3, nanog* and *sox19b*) bind to nucleosomes at promoters and enhancers during blastulation, leading to recruitment of additional TFs, chromatin remodelling enzymes, and polymerases (Zhou and Heald et al., 2023; Barral and Zaret, 2024). Expression profiles of these pioneer factors peak at early developmental stages (usually early blastula) with their expression plummeting as development progresses. As pioneer factors find their target sequences on nucleosomes, histones are also marked and redistributed across the genome (Sinha et al., 2023). For example, writer (*brd4*) and readers (*ep300a* and *ep300b*) for H3K27ac (Chan et al., 2019) steadily increase expression until the shield stage. H3K4me3 writer (*kmt2b*) and reader genes (*taf3*) show similar expression patterns (Hyun et al., 2017)

### Gastrulation, cell fate definition and asynchronous response

As the embryo approaches late blastulation, different cells undergo different paths of differentiation and migration, eventually giving rise to different cell types with committed fates and functions (Veil et al.,2019, Pálfy et al., 2020). Consistent with this information, late blastula seems to be the starting point for the ATAC-Seq and ChIP-Seq assay.

The late blastula to shield was the second most dynamic transition in terms of differential gene expression. Most enriched GO terms were consistent with the convergent extension during gastrulation and the start of pattern specification that happens in the late blastula to shield transition (Wallingford et al., 2002; Fulton et al., 2020). Moreover, the first enriched GO terms associated with morphogenesis were associated with this stage, as discrete cell populations show-up along the blastula stage. Expression of marker genes associated with the extra embryonic enveloping layer and yolk-syncytial layer, such as *krt18a.1* and *gata3* (Xu et al., 2024), were detected in our study from the shield stage onwards. We detected a similar pattern for various members of the Wnt gene family (*wnt5b*, *wnt11*), involved in cell fate determination through the Wnt/β-catening pathway and body axis planning through the Wnt/PCP pathway (Yang and Mlodzkik, 2016; Liu et al., 2022). Finally, we also detected an expression peak during the turbot shield stage for *aldh1a2*, *rdh10*, involved in the synthesis of retinoic acid (RA) as well as genes from the Rar family of RA receptors. RA regulates some key genes involved in patterning like the Hox, Meis and Sox gene families. Recent findings suggest that RA signalling may rewire the chromatin landscape leading to CRE activation, and exogenous RA induces chromatin binding of HOXB1B, MEIS2B and SOX3 TFs (Moreno-Oñate et al., 2024).

Cell fate is driven by interactions between TFs and CREs with accessible chromatin, allowing TF binding (Mircea and Semrau, 2021). However, the number of DARs in gene promoters between consecutive stages was lower than the number of DEGs, with a small, non-significant overlap according to the hypergeometric test. This suggests that most promoters are already accessible without major alterations of global chromatin accessibility between consecutive stages. This, asynchronism between chromatin accessibility of CREs and gene expression has been shown previously (Ma et al., 2020; Wike et al., 2021; Lin et al., 2023). Our data add weight to this notion, identifying chromatin state dynamics for 29 DEGs expressed throughout embryogenesis. Each show accessible promoter regions from the late blastula stage, irrespective of further changes in chromatin accessibility or chromatin state complexity as development progresses.

Interestingly, a previous study in zebrafish suggests that promoter ATAC peaks generally precede transcriptional activity, but in contrast, the majority of distal CREs (which include enhancers) become accessible after gene expression starts (Liu et al., 2024). This past work fits well with our findings, where 20.86% of the DARs across development overlapped enhancer regions, compared to 12.64% for promoter regions.

### Segmentation and organogenesis

By the end of gastrulation, the embryo starts a sequential patterning process along its axis, laying the groundwork for the body plan and organogenesis (Schröter et al., 2008). Accordingly, we detected enriched terms relating to somitogenesis and the development of various tissues and organs were found in mid-to-late development (Kimmel et al., 1995; Schmidt et al., 2013; Cheng et al., 2023) and in the transitions between consecutive stages across segmentation (Miao and Porquié, 2024), while terms associated with locomotion and behaviour were enriched later in the prehatch stage (Pohl, 2019). During the shield to early segmentation transition and subsequent stages, we observed the expected expression changes of candidate master genes related to patterning (*bmp2b*, *bmp4*), chondrogenesis (*foxa2*, *foxa3*) osteoblastogenesis and osteoblast differentiation (*sp7*, *dlx3b*, *dlx5a*), muscle development (*dag1*, *myod1*) and nervous system development (*meis2b*, *sox2*, *neurog1*), among others (Machon et al., 2015; Hamed et al., 2022; Petratou et al., 2024).The diversity and complexity of developmental and regulatory pathways at the onset of somitogenesis is reflected in the number of DEGs and DARS during the shield to early segmentation transition.

Interestingly, the number of DEGs steadily decreased as segmentation progressed, while the number of DARs dramatically decreased in each transitions after early segmentation, most notably from mid-to-late segmentation. This may result in comparatively reduced transcriptional dynamics once the body plan is established, as well as stabilization of the chromatin landscape, with the last major changes in the epigenetic landscape occurring in early segmentation.

As ontogenesis continues after hatching, important biological processes were not detected. Looking at the inactive transcriptome during embryogenesis, we identified enrichment of terms related to immune system and feeding behaviour, a result that aligns with studies on zebrafish, where immune-like response, such as phagocytic activity was not detected until 1-day post-fertilisation (Herbomel et al., 1999; Lange et al., 2023) and adaptive immunity becomes active after three weeks post hatching (Lam et al., 2004; van der Vaart et al., 2012). On the other hand, feeding behaviour is not properly active until complete yolk absorption and the mouth is fully developed (Yúfera and Darías, 2007; Debernardis et al., 2025; Lange et al., 2023).

### Tracing Hox gene evolution and embryogenesis

Hox genes are fundamental in shaping the anterior-posterior body plan and structure of diverse multicellular taxa, as well as governing spinal ossification and regulating organogenesis (Fromental-Ramain et al., 1995; Hubert et al., 2023; Petratou et al., 2024). Moreover, variations in their expression are linked to body structure differences across species, and disruptions in Hox expression can lead to developmental disorders (Cumplido et al., 2024). Comparative analysis of Hox organization and expression can help in better understanding the genetic basis of congenital abnormalities, while also providing evidence of shared ancestry between species.

Hox genes are organized in clusters of up to 13 genes, with a spatial organization that correlates with their temporal and spatial expression patterns (Duboule, 1994). Teleosts underwent a three lineage-specific whole genome duplication events that resulted in up to eight Hox clusters (Wang et al., 2021; Ozernyuk and Schepetov, 2022). However, Hox clusters have been modified by gene loss and co-option during teleost evolution, resulting in significant variation among different lineages (Ozernyuk and Schepetov, 2022). In both turbot and zebrafish, the b paralogs of each initial cluster (HOXAb, HOXBb, HOXCb, HOXDb) have been most affected by this loss, even resulting in the complete loss of some clusters, such as turbot cluster HOXCb and zebrafish HOXDb. Gene loss / downregulation however has not removed any of the 13 Hox genes from the collinear expression chain, as the combined sum of all clusters in each of the species retains at least one Hox gene expressed in anterior (*hox-1* and *hox-2*), antero-central (*hox-3*), central (*hox-4* to *hox-8*) and posterior (*hox-9* to *hox-13*) regions of the embryo.

The majority of orthologous active Hox genes between both species followed a similar expression pattern, increasing from early segmentation and reaching the maximum mRNA levels during later segmentation and pharyngula stages. Although we detected eight Hox orthologs with similar but slightly asynchronous patterns between turbot and zebrafish, we cannot discard that these differences might not be biologically relevant, as both studies were not planned to sample exactly the same developmental stages. However, we found 12 Hox orthologous genes with notably distinct patterns between species. Among them, some early expressed Hox genes in turbot, specially *hoxb8a* and *hoxc5a*, showed relatively high expression at the late blastula stage (∼20 and ∼25 TPM, respectively), while *hoxb1b* showed highest expression in the shield stage (∼75 TPM). To our knowledge there are no reports of Hox expression during blastulation in vertebrates, where the earliest Hox gene expression occurs during gastrulation (Wacker et al., 2004), although there have been reports of Hox gene expression during blastulation in cnidarians (DuBuc et al., 2018).

A significant proportion of turbot Hox paralogs have diverged in their expression pattern, as illustrated by the striking differences observed for *hoxa11*, *hoxa13* and *hoxd9*, or the similar but asynchronous expression patterns for *hoxa2*, *hoxa10* and *hoxb1*. This illustrates that, although paralogous genes can conserve their expression patterns resulting in functional redundancy (Boucherat et al., 2013; Hunter and Prince, 2022), they often diverge, implying a specialised or distinct functional role (McClintock et al., 2001). This can also result in the complete loss of function (Kuraku and Meyer, 2009), as happened with the HOXCb cluster, and seems to be happening with the *hoxa9* and *hoxd4* genes, where paralog b shows much lower levels of expression. It is worth mentioning that even paralogous pairs that have highly similar expression properties may show spatially distinct expression in different cell types or locations in the embryo. Future works that incorporate spatial transcriptomics or single cell technology will build upon the developmental atlas of turbot that has been generated in this study.

## CONCLUSIONS AND PERSPECTIVES

Our study presents the first comprehensive atlas of regulatory elements involved in turbot embryogenesis, contributing to the AQUA-FAANG project under the broader FAANG initiative. This epigenomic atlas offers valuable insights into the molecular mechanisms driving turbot embryogenesis and serves as a unique resource for advancing selective breeding strategies. Future research will benefit from integrating the regulatory annotations identified in this study with genetic variants from whole genome re-sequencing. Such integration will aid in prioritizing causal genetic variants related to growth and reproductive traits.

## MATERIALS AND METHODS

### Embryos

A standard batch of fertilized turbot eggs was provided by Stolt Sea Farm SA (Ribeira, Spain) 2 hours post fertilization (hpf). Eggs were housed in an aerated aquarium with saltwater at the facilities of the Department of Genetics in the Veterinary Faculty of the University of Santiago de Compostela (Spain) at 14.0°C (±0.7°C) until hatching. All samples were collected before hatching (90hpf).

### Protocols

Detailed protocols for egg dechorionation, embryo preservation, RNA isolation and ATAC-Seq and ChIP-Seq procedures (including library preparation) are available in the FAANG repository (data.faang.org; URLs for protocols in **Supplementary tables 1 and 2**).

### Full egg preservation for RNA-Seq

Three pools each of 50 to 150 embryonated eggs were preserved for RNA extraction of 12 developmental stages, with the number of eggs in each pool depending on the sampled stage (**Figure 8**). The desired number of eggs were collected in 2ml safe-lock tubes from the surface of the water with a Pasteur pipette. Saltwater was extracted from the tubes using P1000 and P200 pipettes. Eggs were then resuspended in 1 ml of TRIzol reagent (Invitrogen) and frozen at -80°C for long term storage.

**Figure 8.**
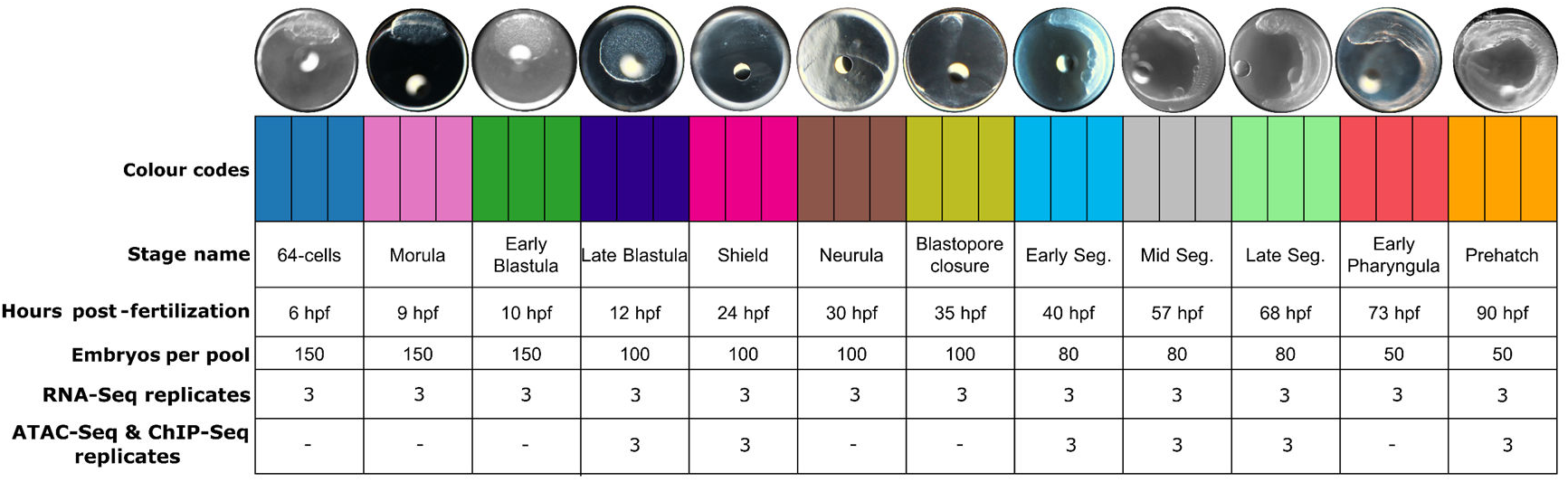
Sampling stages for the transcriptomic and epigenomic assays performed in this study. The colour of each stage is maintained throughout the manuscript figures to refer to each stage.

### RNA isolation and sequencing

After sample thawing, TRIzol was removed and substituted with 700µl of Qiazol Lysis Reagent (miRNeasy Mini Kit, QIAGEN). Two zirconia beads (2.8-3.3 mm) were added to each tube before disrupting the eggs for 2 min, 20Hz in TissueLyser II (QIAGEN). After disruption, total RNA was extracted and purified using the miRNeasy Mini Kit (QIAGEN), following a protocol for “Total RNA extraction for turbot eggs” (**Supplementary table 1**). RNA integrity and quantity were evaluated using a Bioanalyzer (Bonsai Technologies, Madrid, Spain) and NanoDrop® ND-1000 spectrophotometer (NanoDrop® Technologies Inc., Wilmington, DE, USA). The RNA integrity number (RIN) averaged 9.5 across all samples and was always > 7.7. RNA samples were delivered to Novogene (UK) for library preparation using NEBNext Ultra Directional RNA Library Prep Kits for Illumina and sequenced using an Illumina NovaSeq S4 platform to generate 150 bp paired-end reads.

### RNA-Seq data processing

RNA-Seq data was processed using nf-core/rnaseq 3.10.1 (Patel et al., 2023) with default parameters, using the turbot Ensembl genome ASM1334776v1 (Martínez et al., 2021) as reference. In brief, the pipeline evaluated quality of raw reads using FASTQC (Andrews, 2010) and trimmed adapters and low-quality nucleotides using Trim Galore! (Martin et al., 2011). Reads were then mapped to the genome using STAR (Dobin et al., 2013).

### Differential expression analysis

Normalized transcript read counts were obtained using RSEM (Li and Dewey, 2011). After nf-core processing, the resulting count tables were filtered to remove genes with expression below 5 transcripts per million (TPM < 5) and represented in less than two samples across all developmental stages. The genes were clustered in gene expression correlation network using BioLayout (Theocharidis et al., 2009) to explore patterns among expressed genes with the following parameters: 0.94 Pearson correlation threshold; 2.1 cluster granularity, minimum 8 genes per cluster, Markov clustering and Fruchterman-Reingold layout algorithm

### Egg dechorionation for ATAC-Seq and ChIP-Seq procedures

Four Petri dishes, each with 200 eggs per dish, were prepared 3 h before the estimated sampling point for six developmental stages (**Figure 8**). Saltwater was removed from three Petri dishes using a Pasteur pipette, and eggs were resuspended in a pronase solution (from *Streptomices griseus*, ROCHE, 2 mg/ml), leaving the fourth dish as control. All four Petri dishes were transferred to an oscillator (medium velocity) at 19°C (±1°C) for 2.5 to 3.5 h depending on developmental stage, the longer the earlier the stage. The consistency of the chorion was checked by pipetting 10 embryos of each plate after 2.5 h every 20 min. Once the chorion was weakened, the pronase solution was discarded with a Pasteur pipette, and eggs washed twice with PBS. The chorion was then removed by pipetting the eggs up and down, with freed embryos collected in 1.5 LoBind tubes with PBS.

### Cell isolation, library preparation and sequencing for ATAC-Seq

Following a standard protocol for “Cellular isolation of turbot embryos” (**Supplementary table 1**), freshly dechorionated embryos were disaggregated with two distinct procedures, depending on embryo stage. Late-blastula embryos were disaggregated by pipetting up and down 10 times with a P1000 pipette, and vortexed at slow speed for 15 s. Samples for the other five stages were resuspended in 1 ml of Trypsin-EDTA and left under rotation (50 rpm, room temperature -RT-) for 5 min (embryonic shield and early somitogenesis), 10 min (mid and late somitogenesis) or 40 min (prehatch). The trypsin reaction was stopped with 500 µl of cold, heat-inactivated FBS, then centrifuged at 500 g for 6 min at 4°C, and resuspended in 1 ml cold PBS. In total, three replicates for each developmental stage were taken for ATAC-Seq (Figure 1).

The number and integrity of nuclei was assessed with a haemocytometer (minimum of ∼50,000 nuclei in a 16.5 µl suspension; ∼3,000 nuclei/µl) before carrying out the Tn5 transposase reaction with Illumina Tagment DNA TDE1 enzyme (Illumina) using MS-100 Thermo-shaker incubator (Hangzhou Allsheng Instruments) at 37°C, 30 min, 1000 rpm, following the standard “OmniATAC protocol” (Corces et al., 2017; **Supplementary table 2**). The resulting DNA was purified with a MinElute PCR purification kit (Qiagen), and DNA concentration assessed with a Qubit using the dsDNA HS kit (ThermoFisher Scientific). Library amplification was carried out with Integrated DNA Technologies (IDT) for Illumina UD Indexes (96x, Plate A, Set 1, Illumina). Library size selection was performed to remove fragments below 180 bp and above 700 bp using AMPure XP beads (Beckman Coulter). Finally, DNA fragment size distribution was assessed with the Bioanalyzer High Sensitivity DNA Assay kit (Agilent Technologies). ATAC-Seq libraries were delivered to Novogene (UK) and sequenced on an Illumina NovaSeq S4 platform generating 150 bp paired-end reads.

### Embryo cross-linking for ChIP-Seq

Three replicates were taken for each of the six developmental stages (Figure 1). The freshly prepared tubes with pools of dechorionated embryos were centrifuged at 500 g for 6 min at 4°C. After removing the supernatant, embryos were resuspended and incubated in 875 µl of 1% formaldehyde solution on a rotating mixer device (Hula Mixer) at 50 rotations per minute for 15 min at room temperature. Next, 125 µl of 1M glycine solution was added to quench the reaction, incubated under rotation at 40 rotations per minute for 10 min at room temperature. Tubes were centrifuged at 500 g for 6 min at 4°C, removing the supernatant, and resuspended in cold PBS before storage at -80°C.

### Library preparation and sequencing for ChIP-Seq

The frozen crosslinked embryos were thawed on ice. The number and integrity of nuclei was assessed with a haemocytometer (minimum 100,000 nuclei). Nuclei were pelleted and resuspended in complete sonication buffer, and the chromatin sheared using a Covaris S2 focused ultrasonicator with the following parameters: 2% duty cycle, intensity 3, with 200 cycles per burst, at 4°C for 7 min (**Supplementary table 2**).

Due to the expected low number of recovered nuclei, the immunoprecipitation (IP) and library preparation steps were performed following a modified µChIPmentation protocol (Diagenode, **Supplementary table 2**) using Diagenode antibodies for immunoprecipitation (30 ng of chromatin per IP) of three histone marks: H3K4me3 (marking active promoter regions; cat. No. C15410003; 1.3 µg/µl), H3K27ac (marking active enhancer and promoter regions; cat. No. C15410196; 2.8 µg/µl) and H3K27me3 (marking Polycomb repressed regions; cat. No. C15410195; 1.1 µg/µl). Each of the three replicates per developmental stage were subjected to IP. Additionally, for each developmental stage, 4 ul of pooled chromatin (∼1.34 ul per replicate) were set aside to be used as input reference samples, without IP.

Before sequencing, quantity and quality of purified libraries were assessed using the Qubit DNA HS kit (ThermoFisher Scientific) and the High Sensitivity DNA Assay kit (Agilent Technologies), respectively. A minimum of 60% of the chromatin was required to have a size distribution between 200-700 bp (centred around 350-400 bp). ChIP-Seq libraries were delivered to Novogene (UK) for sequencing on an Illumina NovaSeq S4 platform generating 150 bp paired-end reads.

### ATAC-Seq and ChIP-Seq data processing

ATAC-Seq and ChIP-Seq data were processed using the nf-core/atacseq v1.2.1. and nf-core/chipseq v1.2.2 pipelines, respectively (Ewels et al., 2023), run with the narrow_peak option for the ATAC-Seq, and H3K4me3 and H3K27ac ChIP-Seq datasets, and with the broad_peak option for H3K27me3. Other parameters were default. Briefly, quality assessment of the reads was carried out with FASTQC (Andrews, 2010), and adapters and low-quality bases were trimmed with Trim Galore! (Martin et al., 2011). Reads were mapped to the turbot genome using BWA (Li and Durbin, 2009). Further filtering was done with SAMtools (Danecek et al., 2021), BAMtools (Barnett et al., 2011) and Pysam (Danecek et al., 2021). Genome-wide IP enrichment relative to controls was done with deepTools (Ramírez et al., 2014) and broad/narrow peaks were called using MACS2 (Zhang et al., 2008).

An ATAC-Seq consensus peak list was generated using ATAC-Seq peaks detected consistently in two or more biological replicates in at least one stage using the R/Bioconductor package DiffBind (Stark and Brown, 2011) with default settings. Additionally, a previous turbot ChIP-Seq blacklist (Aramburu et al., 2023, of high signal (5.58% of the turbot genome) and low mappability regions (1.40%) was incorporated for downstream filtering.

### Chromatin state inferences

Genome-wide chromatin states were modelled across turbot embryogenesis using ChromHMM (Ernst and Kellis, 2017) integrating the ChIP-Seq and ATAC-Seq data. ChromHMM models including 8 to 15 states were tested, with the aim of maximizing the number of non-redundant, biologically relevant chromatin states (Pan et al., 2021; Baranasic et al., 2022; Vu and Enst, 2022). The ATAC-Seq consensus peakset was intersected and annotated with the chromatin-state map for each developmental stage, and peaks further annotated as representing promoters (-2000 to +600 bp of each gene TSS) or being located in introns, exons or intergenic regions. ATAC-Seq peaks annotated as promoters with active-promoter states, and introns/intergenic regions with active-enhancers states were retrieved.

### Differential gene expression and chromatin accessibility analysis

Differentially expressed genes (DEGs) between consecutive pairs of the six developmental stages were identified using the R/Bioconductor package DESeq2 v1.44 (Love et al., 2014), resulting in five comparisons. Genes with false discovery rate (FDR) adjusted p < 0.05 were considered DEGs. Using the consensus ATAC-Seq peaks, differentially accessible regions (DARs; adjusted p < 0.05) between the same pairs of consecutive developmental stages were identified using the R/Bioconductor package DiffBind (Stark and Brown, 2011), with default settings.

### Gene ontology analysis

In total, 19 transcriptomic profiles were considered to characterize gene expression across development, consisting of: i) the total active transcriptome across the whole turbot development (all genes expressed across studied stages); ii) the total inactive transcriptome (genes < 5 TPM across stages); iii) 12 stage-specific profiles, including genes expressed in each particular stage (see material and methods); and iv) five differential expression profiles between the six consecutive main stages. Gene ontology (GO) enrichment for each profile was performed using ShinyGO v0.80 (Ge et al., 2020) for terms associated with Biological Process (BP), which were ranked by statistical significance (FDR-adjusted p < 0.05). The total active transcriptome was used as background for the GO analyses for the 12 stage-specific (i) and five differentially expressed profiles (ii). All resulting datasets were summarized using Revigo v1.8.1 (Medium complexity, FDR enrichment used as scaling parameter, Supek et al., 2011)

### Transcription factor motif analysis

For each list of DARs annotated as active promoters or enhancers, enriched transcription factor (TF) binding motifs (TFBM) included in the HOMER software (Heinz et al., 2010) were identified using the findMotifsGenome.pl function (settings: -size given -mask-mset vertebrates). Random genomic regions with GC-content matching each input genomic list were used as the background for automatic motif analysis by HOMER. Only TFBM with adjusted *p* < 0.05 after Bonferroni correction and percentage of target sequences > 10% were considered to be enriched.

GO analysis of the TFs predicted to bind enriched TFBM was performed with Metascape (Zhou et al., 2019) using zebrafish (*Danio rerio*) as reference.

### Comparative analyses with zebrafish

Orthology analysis between the predicted proteomes of turbot (assembly ASM1334776v1) and zebrafish (GRCz11) was performed with OrthoFinder 2.5.4 (Emms and Kelly, 2019), using canonical predicted amino acid sequences only. The orthology trees were rooted with the proteomes of three other teleost species: *Cynoglossus semilaevis* (Cse_v1.0), *Oryzias latipes* (ASM223467v1) and *Lepisosteus oculatus* (LepOcu1).

Turbot and zebrafish genes included in single copy orthogroups (i.e. only one orthologous gene in all included species) were retrieved for comparative analysis. Expression profiles were compared between the six main developmental stages studied in turbot (late blastula, embryonic shield, early segmentation, mid segmentation, late segmentation and prehatch) and the six homologous morphological stages studied by White et al. (2017): dome, gastrula shield, 1-4 somites, 14-19 somites, 20-24 somites and long-pec, respectively.

Graph-based comparison of expression patterns across the six main developmental stages of both species was accomplished with R by grouping the genes in our dataset by their stage of maximum expression. Then, intra-species normalization of the TPM counts was carried out against the top value across all stages considered, within each species. The Spearman correlation coefficient was used to calculate a distance matrix by hierarchical clustering (Complete linkage). Additionally, special attention was paid to the comparison of expression patterns for Hox gene family orthologs between both species, as well as the comparison of expression patterns and levels of expression between the turbot Hox gene paralogs.

## DATA AVAILABILITY STATEMENT

All raw RNA-Seq, ATAC-Seq and ChIP-Seq datasets can be accessed through the ENA repository under accession numbers PRJEB47933, PRJEB47934 and PRJEB57784, respectively. The experimental protocols are summarized in the material and methods section, but in-depth protocols can be accessed through the URLs to the FAANG repository, found in supplementary tables 1 and 2. All gene and protein names used in the manuscript adhere to their official symbols, and supplementary tables provide the turbot-specific ensembl IDs of these genes.

## AUTHOR CONTRIBUTIONS

**OA:** Methodology, Software, Formal analysis, Investigation, Data Curation, Visualization, Writing original draft, Writing – review & editing; **BGP:** Methodology, Investigation, Resources, Writing review & editing; **AJG:** Formal Analysis, Methodology, Supervision, Resources, Writing – review & editing; **ABH:** Software, Formal Analysis, Data Curation, Visualization; **DM**: Funding Acquisition, Methodology, Supervision, Resources, Writing – review & editing; **CB**: Conceptualization, Formal Analysis, Resources, Writing – review & editing, Project Administration, Supervision; **PM**: Conceptualization, Methodology, Formal Analysis, Resources, Writing – review & editing, Project Administration, Supervision.

## ACKNOWLEDGMENTS

We acknowledge the technical support and informatic resources provided by the Centro de Supercomputación de Galicia (CESGA).

